# Organ specificity and commonality of epigenetic aging in low- and high-running capacity rats

**DOI:** 10.1101/2024.04.21.590009

**Authors:** Takuji Kawamura, Csaba Kerepesi, Ferenc Torma, Zoltan Bori, Lei Zhou, Peter Bakonyi, Attila Kolonics, Laszlo Balogh, Mitsuru Higuchi, Lauren Gerard Koch, Steven Loyal Britton, Erika Koltai, Zsolt Radak

## Abstract

**Background:** Epigenetic drift, which are gradual age-related changes in DNA methylation patterns, plays a significant role in aging and age-related diseases. However, the relationship between exercise, epigenetics, and aging and the molecular mechanisms underlying their interactions are poorly understood. The aim of this study was to investigate the relationship between cardiorespiratory fitness (CRF), epigenetic aging, and promoter methylation of individual genes across multiple organs in selectively bred low- and high-capacity runner (LCR and HCR) aged rats.

**Methods:** In this study, we performed reduced representation bisulfite sequencing (RRBS) on LCR and HCR aged rats, focusing on the hippocampus, heart, soleus muscle, and large intestine samples. We calculated various epigenetic indicators, including rat epigenetic clocks, global mean DNA methylation (GMM), mean methylation entropy (MME), and gene-specific promoter methylation.

**Results:** Epigenetic clocks, trained on available rat blood-derived RRBS data did not reflect differences in CRF between LCR and HCR rats across all four organs. However, we observed organ specific differences in GMM and MME between LCR and HCR rats. The direction of these differences was the opposite compared to the age-related changes in the rat blood suggesting that a high CRF may mitigate age-related epigenetic changes in an organ-specific manner. Notably the soleus muscle, exhibited the most pronounced differences in promoter methylation due to CRF. We also identified seven genes whose promoter methylation was consistently influenced by CRF in all four organs. Moreover, we found that age acceleration of the soleus muscle was significantly higher compared to the heart and the hippocampus, and significantly lower compared to the large intestine. Finally, we found that the age acceleration was not consistent across organs.

**Conclusions:** Our data suggest that CRF associates with epigenetic aging in an organ-specific manner and regulates the promoter methylation of individual genes in an organ-specific and organ-common manner. Our findings provide important insights into the biology of aging and emphasize the need to validate rejuvenation strategies in the context of the organ-specific nature of epigenetic aging.

## 1. Introduction

Aging is a complex biological process characterized by a gradual decline in physiological integrity, which leads to impaired function and increased vulnerability to death^1^. The aging rate is influenced by various genetic and environmental factors. Among these factors, interest in epigenetics as an important regulator of aging has been increasing. Epigenetic modifications, including DNA methylation, histone modifications, and non-coding RNA regulation, play a pivotal role in regulating gene expression patterns and consequently contribute significantly to the aging process. Notably, alterations in DNA methylation patterns during aging, termed epigenetic drift, occur across various tissues and cells, and contribute to the progressive dysregulation of gene expression. Therefore, clinical trials have begun to evaluate the efficacy of rejuvenation strategies targeting epigenetic drift, such as lifestyle interventions, plasmapheresis, and the utilization of drugs and dietary supplements^2^.

The beneficial effects of exercise on healthy aging are well documented. In particular, cardiorespiratory fitness (CRF), represented by maximal oxygen uptake (VO_2max_), is an indicator of longevity^3,4^. However, understanding the relationship among exercise, epigenetics, and aging is a complex endeavor, and the molecular mechanisms underlying this interaction remain an ongoing scientific pursuit. Previous research has demonstrated the influence of acute and regular exercise on global and gene-specific promoter methylation, primarily in peripheral blood and skeletal muscle^5,6^. Recent findings, including those of our study, also indicate a negative relationship between physical fitness (including activity levels and CRF) and epigenetic aging based on age-related changes in DNA methylation levels at CpG sites^7–10^. These lines of evidence suggest that exercise exerts a rejuvenating effect on epigenetic aging and has a favorable effect on the aging process. Importantly, the health-promoting effects of maintaining and improving CRF extend to multiple organs and the whole body^11^. However, only one study has investigated the relationship between CRF and epigenetic aging across multiple organs^12^. To the best our knowledge, no study has explored the organ specificity and commonality of promoter methylation in individual genes according to differences in CRF.

Based on extensive epidemiological studies that have shown that VO_2max_ is a strong predictor of morbidity and mortality^3,4^, Koch and Britton formulated the ‘aerobic hypothesis’ that variation in the capacity for oxygen metabolism is the central mechanistic determinant of the difference between complex diseases and health^13^. To test this hypothesis, low- and high-running capacity strains were established by divergent selection breeding for running capacity using a founder population of a genetically heterogeneous N:NIH rat stock (*i.e.*, eight inbred lines: ACI, BN, BUF, F344, M520, MR, WKY, WN)^14,15^. Notably, VO_2max_, measured throughout adulthood, was a reliable predictor of lifespan, and the median lifespan was extended by 8–10 months in high-capacity runners (HCR) than in low-capacity runners (LCR) without extending maximum lifespan^15^. This finding implies that the LCR rats were more susceptible to disease, supporting the healthspan-extending effects (not lifespan-extending effects) of exercise. Therefore, this rat strain is ideal for elucidating the unknown mechanisms underlying the correlation between VO_2max_ and healthspan, and how VO_2max_ regulates epigenetic aging and promoter methylation, which are associated with morbidity and mortality, across multiple organs.

In this study, we performed reduced representation bisulfite sequencing on the hippocampus, heart, soleus muscle, and large intestine of selectively bred LCR and HCR aged rats, which are known for their different disease risks and life expectancy^15,16^. Our aim was to elucidate the relationship between CRF, epigenetic aging, and promoter methylation of individual genes across multiple organs in this rat model.

## 2. Results

### 2.1. Global changes in the rat methylome during aging is delayed in HCR compared to LCR rats

To determine the relationship between differences in CRF, epigenetic age, and global changes in the methylome with aging across multiple organs, we obtained DNA methylome data from the hippocampus, heart, soleus muscle, and large intestine of 16 (LCR: *n* =8 and HCR: *n* =8) 23–24 months old rats, which were selectively bred for 44 generations based on endurance running capacity^13^ (**Fig. 1A and Table 1**). At least 48 h before organ sample collection, the maximum treadmill running capacity of LCR and HCR rats was measured, which revealed that VO_2max_ total running time and total running distance were higher in the HCR group than in the LCR group. These results indicated that CRF, which gradually declines with age, retains differences in these rat models even at this age (**Fig. 1B**). Genomic DNA extracted from each organ sample was subjected to RRBS, and we obtained data for > 6 million CpG sites in the 64 samples.

**Figure 1.**
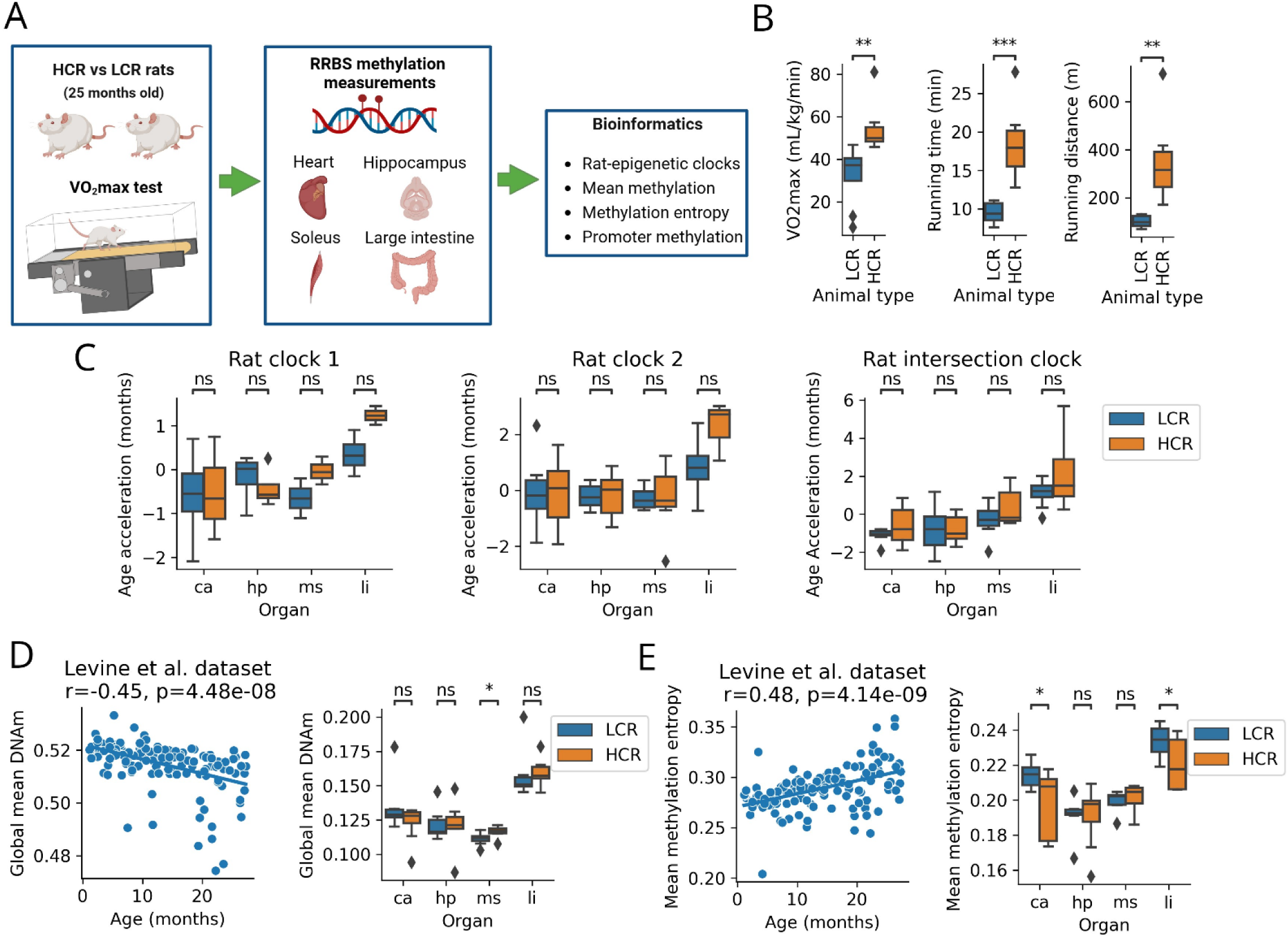
Global changes in the rat methylome during aging is delayed in HCR rats compared to LCR rats. **A.** The overview of the study design. **B.** maximal treadmill running capacity of LCR and HCR rats. **C.** Epigenetic age accelerations of three rat clocks in multiple organs of LCR and HCR rats. **D.** Global DNAm levels of 1 028 452 CpG sites during aging (using Levine rat blood dataset) and their comparison in multiple organs of LCR and HCR rats. **E.** Mean methylation entropy of CpG sites during aging (using Levine rat blood dataset) and their comparison in multiple organs of LCR and HCR rats. LCR: low-capacity runner, HCR; high-capacity runner, ca: heart, hp: hippocampus, ms: soleus muscle, li: large intestine. *r*: Peason’s correlation coefficient, *p*: *p* value of each test, *: 0.01 < *p* ≤ 0.05, **: 0.001 < *p* ≤ 0.01, ***: *p* ≤ 0.001, ns: not significant.

**Table 1.** Chronological age, body and organ weight of LCR and HCR rats.

| | LCR ($n = 8$) | HCR ($n = 8$) | $p$ value |
| --- | --- | --- | --- |
| Age (months) | 25.3 ± 0.3 | 25.4 ± 0.4 | 0.888 |
| Body weight (g) | 312 ± 40 | 277 ± 50 | 0.154 |
| Hippocampus (mg) | 59 ± 5 | 58 ± 4 | 0.430 |
| Heart (mg) | 933 ± 79 | 926 ± 53 | 0.830 |
| Soleus (mg) | 99 ± 16 | 108 ± 16 | 0.297 |
| Large intestine | N.A. | N.A. | - |

Age indicates the chronological age at the time of conducting the dissection. Organ weights are not shown for the large intestine, as part of it was cut off and stored. LCR: low-capacity runner, HCR; high-capacity runner, N.A: not applicable.

To develop a RRBS-based rat epigenetic clock of our rat data, we downloaded the processed methylation files (CGMap) of the GSE161141 dataset^17^ and developed ‘Rat clock 1’ and ‘Rat clock 2,’ respectively (see details in the Materials and methods section). Both clocks were trained on 80 % (*n* = 107) of the random samples using the machine-learning tool glmnet and tested on 20 % of the samples (*n* = 27). The performance on the test set was MAE = 4.3 months and *r* = 0.88 by Rat clock 1 and MAE = 4.4 months and *r* = 0.87 by Rat clock 2. A total of 39 CpG sites were used by Rat clock 1, and 27 CpG sites were used by Rat clock 2. Age acceleration was calculated from the residuals of the regression line on the test set for the two rat clocks. Owing to the random nature of RRBS, the number of clock CpG sites in the application dataset may be limited. Therefore, we found the common CpG site in the training dataset (*i.e.*, Levine’s rat data) and the samples of the application dataset (*i.e.*, our rat data) and developed a ‘Rat intersection clock,’ on a common site according to the intersection clock method^18^. As we applied these three rat clocks to our data, we unexpectedly observed no difference between the LCR and HCR groups in the acceleration of the age of the rat clocks in any organ (**Fig. 1C**).

We also used the Levine rat blood dataset to test the association between the GMM, MME, and aging. We found that GMM was negatively correlated (*r* = −0.45; *p* = 4.48e-08) (**Fig. 1D**) and MME was positively correlated (*r* = 0.48; *p* = 4.14e-09) (**Fig. 1E**) with age. We then compared the GMM and MME in each organ between the LCR and HCR groups. GMM in the LCR soleus muscle samples was lower than that in the HCR samples (**Fig. 1D**). The MME was higher in the LCR heart and large intestine samples than in the HCR samples (**Fig. 1E**).

Altogether, while the applied blood-based epigenetic clocks did not show age acceleration differences between LCR and HCR rats for any of the examined organs, we observed that the HCR rat methylome as a whole expressed a younger state compared to the LCR rat methylome.

### 2.2. LCR and HCR rat had the top hit genes in the soleus muscle and seven common genes across all four organs

To identify the best-hit genes and common genes affected by CRF-induced promoter methylation across all four organs, we calculated the mean methylation levels of CpG sites in the promoter region of each sample. Among the 14 366 examined genes, we identified the six best-hit genes with the lowest *p*-values across all four organs in the LCR and HCR groups. These best-hit genes were *Acot5-ps1* (Acyl-CoA Thioesterase 5 Pseudogene 1), *LOC102547081* (uncharacterized gene), *Stk24* (Serine/Threonine Kinase 24), *Tuba4a* (Tubulin Alpha 4a), *Sfmbt2* (Scm-Like with Four Mbt Domains 2), and *LOC100359655* (uncharacterized gene), all of which were observed in the soleus muscle with an uncorrected *p*-value lower than 3.208E-06 and Bonferroni corrected *p* < 0.0461 (**Fig. 2A**). Pseudogenes, such as *Acot5-ps1*, do not typically have a functional role and are non-coding DNA sequences. *Stk24* is a serine/threonine kinase involved in signal transduction pathways that influence cellular processes associated with growth, differentiation, and various physiological functions. *Tuba4a* encodes the tubulin protein, which is a structural component of microtubules. Microtubules are essential for various cellular processes including cell division, intracellular transport, and cell shape maintenance. *Sfmbt2* is associated with epigenetic regulation, involved in the maintenance of chromatin structure, and plays a role in controlling gene expression and cellular development. *LOC102547081* and *LOC100359655* are uncharacterized genes and their precise physiological roles remain unknown.

**Figure 2.**
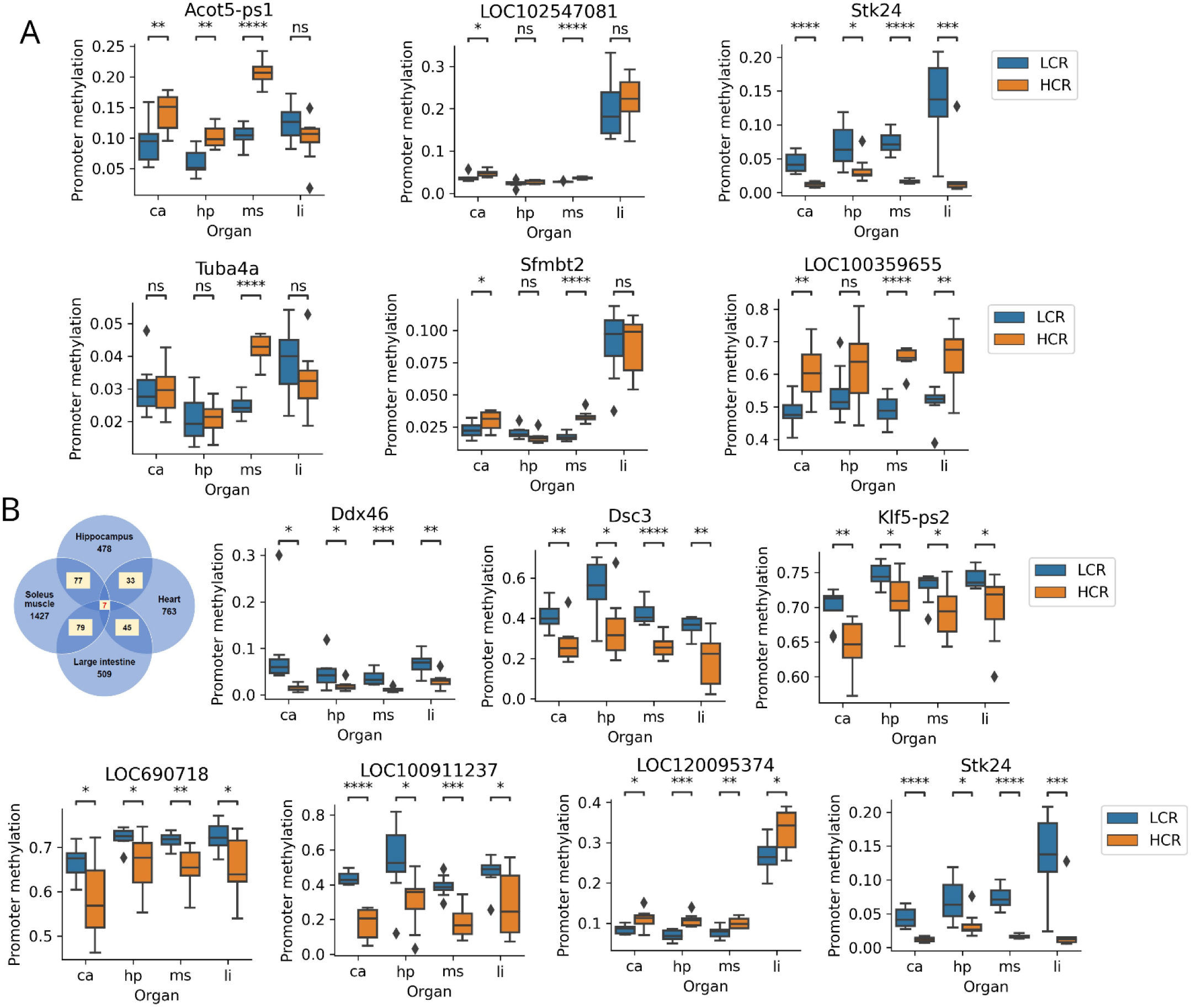
Organ commonality of promoter methylation in individual genes in LCR and HCR rats. **A.** Six best hit genes across the four targeted organs of LCR and HCR rats. **B.** The seven common genes influenced in promoter methylation by CRF across four organs. LCR: low-capacity runner, HCR; high-capacity runner, ca: heart, hp: hippocampus, ms: soleus muscle, li: large intestine. *Acot5-ps1*: Acyl-CoA Thioesterase 5 Pseudogene 1, *Stk24*: Serine/Threonine Kinase 24, *Tuba4a*: Tubulin Alpha 4a, *Sfmbt2*: Scm-Like with Four Mbt Domains 2, *LOC102547081/LOC100359655*: uncharacterized genes. *Ddx46*: DEAD-Box Helicase 46, *Dsc3*: Desmocollin-3, *Klf5-ps2*: Kruppel-Like Factor 5 Pseudogene 2, *LOC690718/LOC100911237/LOC120095374*: uncharacterized genes. *: 0.01 < *p* ≤ 0.05, **: 0.001 < *p* ≤ 0.01, ***: *p* ≤ 0.001, ****: *p* ≤ 0.0001, ns: not significant.

We also identified a distinct set of seven genes that exhibited consistent differential methylation levels across all the four organs examined in the LCR and HCR groups (**Fig. 2B**). The seven identified common genes comprised *Ddx46* (DEAD-Box Helicase 46), *Dsc3* (Desmocollin-3), *Klf5-ps2* (Kruppel-Like Factor 5 Pseudogene 2), *LOC690718* (uncharacterized gene), *LOC100911237* (uncharacterized gene), *LOC120095374* (uncharacterized gene), and *Stk24* (Serine/Threonine Kinase 24), and except for *LOC120095374*, hypomethylation was observed in the HCR group when compared to the LCR group across all four organs (**Fig. 2B**). *Ddx46* is an RNA helicase involved in RNA processing and translational regulation. *Dsc3* is a component of desmosomes that plays a critical role in cell-cell adhesion, contributing to tissue integrity and cohesion. *Klf5-ps2* is a transcription factor implicated in the regulation of cell proliferation, differentiation, and growth. *Stk24* is a serine/threonine kinase associated with various cellular functions including signal transduction and cell growth regulation. The precise physiological roles of LOC690718, LOC100911237, and LOC120095374 are yet to be elucidated. Therefore, further investigation is required to determine the specific functions of these genes.

### 2.3. LCR and HCR rat had different methylation levels in gene-specific promoter regions depending on organs

Next, we identified genes characteristic of each organ with different methylation levels in their promoter regions. We used the data on the mean methylation level at the CpG site of the promoter region of each sample, calculated by the method described above, and illustrated the six best-hit genes for each organ with the lowest *p*-value in the comparison between the LCR and HCR groups (**Fig. 3A–D**).

**Figure 3.**
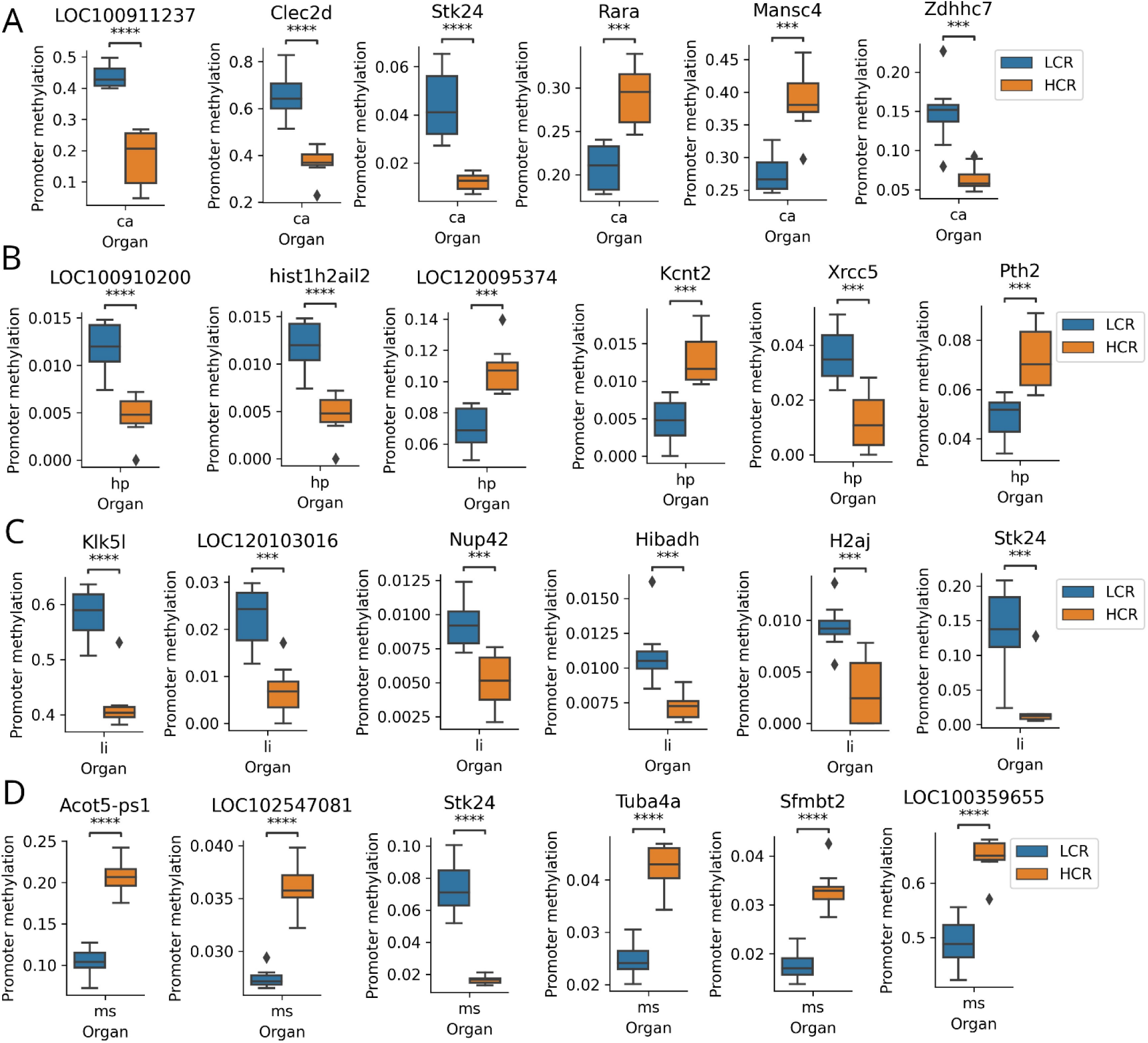
Organ specificity of promoter methylation in individual genes in LCR and HCR rats. Six best hit genes influenced in promoter methylation in the **A.** Heart, **B.** Hippocampus, **C.** Large intestine and **D.** Soleus muscle. LCR: low-capacity runner, HCR; high-capacity runner, ca: heart, hp: hippocampus, ms: soleus muscle, li: large intestine. *Clec2d*: C-type lectin domain family 2, member D, *Stk24:* Serine/Threonine Kinase 24, *Rara*: Retinoic Acid Receptor Alpha, *Mansc4: MANSC Domain Containing 4*, *Zdhhc7*: Zinc Finger DHHC-Type Containing 7, *hist1h2ail2*: Histone Cluster 1 H2A Family Member L2, *Kcnt2*: Potassium Channel, Sodium-Activated, Subfamily T, Member 2, *Xrcc5*: X-Ray Repair Cross Complementing 5, *Pth2*: Parathyroid Hormone 2, *Klk5l*: Kallikrein-Related Peptidase 5-Like, *Nup42*: Nucleoporin 42, *Hibadh*: 3-Hydroxyisobutyrate Dehydrogenase, *H2aj*: Histone H2AJ, *Acot5-ps1*: Acyl-CoA Thioesterase 5 Pseudogene 1, *Tuba4a*: Tubulin Alpha 4a, *Sfmbt2*: Scm-Like with Four Mbt Domains 2, *LOC100911237*/*LOC100910200*/LOC120095374/*LOC120103016/LOC102547081/LOC100359655*: uncharacterized genes. ***: *p* ≤ 0.001, ****: *p* ≤ 0.0001.

In the heart, the best-hit genes were *LOC100911237* (uncharacterized gene), *Clec2d* (C-type lectin domain family 2, member D), *Stk24* (Serine/Threonine Kinase 24), *Rara* (Retinoic Acid Receptor Alpha), *Mansc4 (Mansc4)*, and *Zdhhc7* (Zinc Finger DHHC-Type containing 7) (**Fig. 3A**). *Clec2d* is associated with the immune system and possibly plays a role in immune responses and cell-cell interactions, particularly in the context of immunity and defense against pathogens. *Stk24* is a serine/threonine kinase that is involved in signal transduction pathways, influences cellular processes related to growth and differentiation, and potentially plays a role in various physiological functions. *Rara* is a nuclear receptor that responds to retinoic acid, a form of vitamin A, which plays a crucial role in regulating gene expression and is involved in processes such as cell differentiation and development. Information regarding the specific physiological role of *Mansc4* is limited, and further research is required to elucidate its function. *Zdhhc7* encodes a protein with a DHHC domain potentially involved in palmitoylation, which can affect the function and localization of various proteins, including those involved in cell signaling.

The best-hit genes in the hippocampus were *LOC100910200* (uncharacterized gene), *hist1h2ail2* (Histone Cluster 1 H2A Family Member L2), *LOC120095374* (uncharacterized gene), *Kcnt2* (Potassium Channel, Sodium-Activated, Subfamily T, Member 2), *Xrcc5* (X-Ray Repair Cross Complementing 5), and *Pth2* (Parathyroid Hormone 2) (**Fig. 3B**). The *hist1h2ail2* protein is involved in DNA packaging and gene regulation and is possibly associated with chromatin structure and epigenetic regulation. *Kcnt2* encodes a sodium-activated potassium channel that regulates the electrical activity of cells and is involved in ion transport and cellular excitability. *Xrcc5* is involved in DNA repair and associated with the maintenance of genomic stability and repair of DNA damage caused by various factors. *Pth2* encodes a parathyroid hormone that plays a role in calcium and phosphate homeostasis, thereby influencing bone health and mineral metabolism.

In the large intestine, the best-hit genes were *Klk5l* (Kallikrein-Related Peptidase 5-Like), *LOC120103016* (uncharacterized gene), *Nup42* (Nucleoporin 42), *Hibadh* (3-Hydroxyisobutyrate Dehydrogenase), *H2aj* (Histone H2AJ), and *Stk24* (Serine/Threonine Kinase 24) (**Fig. 3C**). *Klk5l* is possibly a member of the kallikrein-related peptidase family, which includes enzymes involved in various physiological processes such as tissue remodeling, inflammation, and blood pressure regulation. *Nup42* is a nucleoporin component of the nuclear pore complex that regulates the transport of molecules between the nucleus and cytoplasm. *Hibadh* is involved in fatty acid metabolism and breakdown of 3-hydroxyisobutyrate, a metabolite associated with valine catabolism and energy production. The *H2aj* protein is involved in DNA packaging and gene regulation and may play a role in chromatin structure and epigenetic regulation. *Stk24* is a serine/threonine kinase involved in signal transduction pathways that influence cellular processes associated with growth, differentiation, and various physiological functions.

The best hit genes in soleus muscle were *Acot5-ps1*, *LOC102547081 Stk24*, *Tuba4a*, *Sfmbt2*, and *LOC100359655* (**Fig. 3D**), which have been described above.

### 2.4. Age acceleration of the soleus muscle was significantly higher compared to the heart and the hippocampus, and significantly lower compared to the large intestine

In addition to the comparisons between the LCR and HCR groups, we compared the rates of epigenetic aging between organs using data from 16 samples from both the LCR and HCR groups. Age accelerations according to the Rat clock 1 and Rat clock 2 clocks showed similar results and were significantly higher in the large intestine than in the heart, hippocampus, and soleus muscle (**Fig. 4A and 4B**). Using the Rat intersection clock, that is expected to be the most reliable method, the age acceleration of the soleus muscle were significantly higher compared to the heart and the hippocampus, and significantly lower compared to the large intestine (**Fig. 4A and 4B**). GMM was significantly higher in the large intestine than in the heart, hippocampus, and soleus muscle, and was significantly higher in the heart than in the soleus muscle (**Fig. 4C**). MME was significantly higher in the large intestine than in the heart, hippocampus, and soleus muscle, and was significantly higher in the heart and soleus muscle than in the hippocampus (**Fig. 4C**). Overall, the set of epigenetic aging measurements in the large intestine and soleus muscle showed different signatures compared to the other two organs.

**Figure 4.**
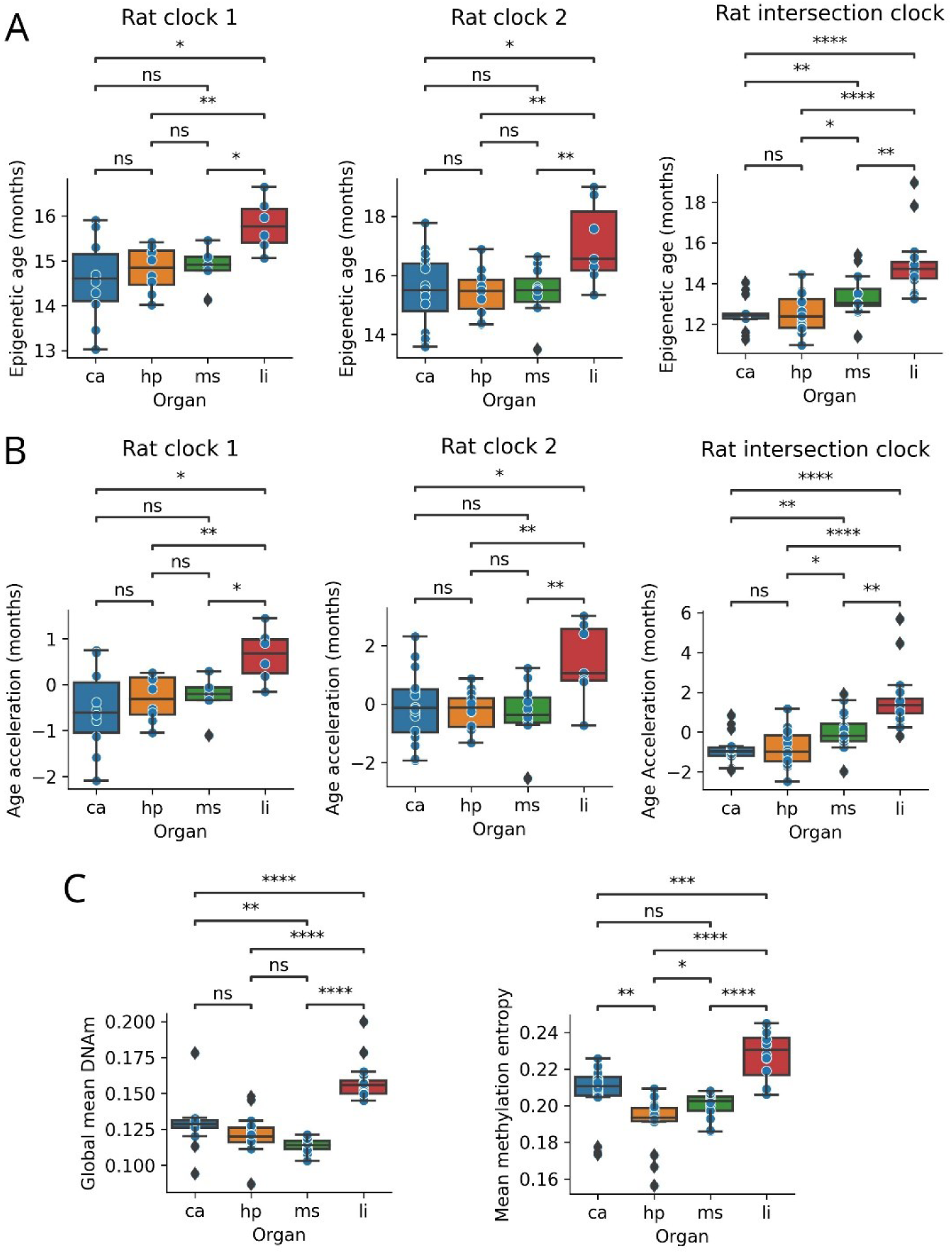
Multi-organ comparisons of epigenetic aging in both LCR and HCR rats. **A.** Multi-organ comparisons of epigenetic age of three rat clocks in both LCR and HCR rats. **B.** Multi-organ comparisons of epigenetic age acceleration of three rat clocks in both LCR and HCR rats. **C.** Multi-organ comparisons of global mean DNAm level and mean methylation entropy in both LCR and HCR rats. LCR: low-capacity runner, HCR; high-capacity runner, ca: heart, hp: hippocampus, ms: soleus muscle, li: large intestine. *: 0.01 < *p* ≤ 0.05, **: 0.001 < *p* ≤ 0.01, ***: *p* ≤ 0.001, ****: *p* ≤ 0.0001, ns: not significant.

### 2.5. Inconsistency of epigenetic age acceleration across organs within in the same individual

As we measured the epigenome of four organs per individual, we had the unique opportunity to investigate whether accelerated (or decelerated) aging is a global process affected to the whole body or rather appear only locally for a specific organ. We found no significant correlations between any organ pair meaning that epigenetic age acceleration is not consistent across organs (**Fig.S2**).

## 3. Discussion

Epigenetic clocks are a promising aging biomarker that can estimate biological age across organs based on age-related changes in DNA methylation patterns, and have demonstrated high predictive power for age-related disease onset and mortality in humans^19,20^. Epigenetic clocks have also been developed in rodents such as mice^21–24^ and rats^17,25^ and pan-species clocks have been proposed, including human-rat pan-tissue clock^26,27^. These findings and latest studies^28^ suggest that age-related changes in DNA methylation patterns are a common mechanism of aging across mammals and, simultaneously, provide a useful tool for evaluating the effectiveness of intervention strategies to delay aging in a relatively short timescale. Epigenetic age can be delayed by caloric restriction, diet quality, growth hormone receptor knockout, plasmapheresis, and lifestyle interventions in rodents and human^21–24, 29–34^. Cross-sectional studies in human populations have shown that physical activity and CRF are associated with delayed epigenetic age progression^7–10^. In mice, late-life exercise training may delay skeletal muscle epigenetic aging via skeletal muscle-specific epigenetic clock(s)^35,36^. In contrast, we demonstrated that blood-based rat epigenetic clocks do not consistently reflect differences in CRF across all organs. This result may be partly explained with that the rat clocks were based on whole blood samples and that our dataset consisted of organ samples. However, in previously studies, blood-based RRBS clocks successfully captured rejuvenation effects in other tissues and cells^21,34,37^. Another possible limitation of our analysis, that the training set of the rat clocks contained only males while the set contained only females. This may influence the accuracy of the clock and explain the lower predicted age compared to the chronological age (**Fig. 4A**). The tissue-specific analyses of another study using the same rat model but the mammalian methylation array platform revealed a lower epigenetic age (adjusted for chronologic age, generation, sex and treadmill running distance at 3 months of age) in HCR rats for adipose, skeletal muscle, cardiac muscle (pan-tissue rat clock; *p <* 0.04), and liver (human-rat clock for relative and chronologic age; *p* < 0.01) compared to LCR rats^12^. However, they did not examine hippocampus, soleus muscle, and large intestine. The study also demonstrated that the effect of CRF on epigenetic age was strongest in young rats compared to old rats; therefore, we cannot exclude the possibility that the effect of CRF may be reduced in old rats^12^. Lastly, the study measured male and female rats, while we measured only females. It is possible, that the effect of high CRF on the rate of aging is milder in females compared to males. This would be consistent with our recent study where we found that standard blood test-based age acceleration largely decreased in male athletes compared to the healthy male controls but did not significantly decrease in female athletes compared to healthy female controls^38^. Other previous study has shown that the age acceleration in the high-fit group was significantly lower than in the medium-to low-fit group for both men and women, but the difference was slightly lower for women^8^. Our previous study also found a significant negative relationship between CRF and age acceleration, but this study included only men^9^. Most other previous studies have examined the relationship between physical activity and age acceleration in the same cohort of men and women (adjusted for sex) through cross-sectional designs^9^. Considering the overall findings of previous studies that suggest that exercise may delay epigenetic age progression, the relationship between exercise and multi-organ epigenetic age is worth investigating, along with the development of a rat clock that can capture biological aging with high precision across multiple organs, as well as their sex differences. In addition, future comparisons of epigenetic aging in LCR/HCR rats at several life stages will be needed to elucidate the relationship between CRF and trajectories of epigenetic aging.

Measurements that may capture changes in DNA methylation during aging include GMM and MME. Although evidence suggests that GMM generally declines with age, the relationship between these factors appears to vary among organs and measurement methods^39^. Results from blood-based methylation assays indicated a negative correlation between GMM and age in mice^40^; however, this relationship was unknown in rats. In this study, we showed for the first time in rats that a negative correlation exists between blood GMM and age and that high CRF mitigates the age-related decline in GMM only in the rat soleus muscle. Entropy is a measure of disorder in an aging system, in which death is the maximum disorder^41^. MME has been shown to increase during aging in mice, naked mole-rats, and humans^25,40,42^; however, the relationship between the MME and aging in rats remains unknown. We found a positive correlation between MME and age, and high CRF mitigated age-related increases in MME in the rat heart and large intestine. These findings on GMM and MME suggest that maintaining high CRF levels may delay age-related changes in DNA methylation patterns in an organ-specific manner. A previous study showed that acute exercise decreases global promoter methylation in human skeletal muscles^5^. Chronic exercise also affects global promoter methylation in human skeletal muscles^5^. However, no study has investigated the relationship between the GMM, MME, and CRF in multiple organs. Considering these facts, the results of this study may provide an insight into the mechanism of the systemic ‘geroprotective’ effect of exercise. However, GMM and MME are not direct measures of epigenetic age, and the differences between the LCR and HCR groups were organ-specific and inconsistent between GMM and MME, that is, there were group differences in the soleus muscle for GMM, while in the heart and large intestine for MME. Therefore, further studies on the relationship between epigenetic aging and CRF are needed to clarify their organ specificity and organ commonality.

In this study, we performed comprehensive promoter analyses of individual genes, which were not limited to ‘exercise-related genes’ such as peroxisome proliferator-activated receptor gamma, coactivator 1 α (*PGC-1α*), mitochondrial transcriptional factor A (*TFAM*) and peroxisome proliferator-activated receptor δ (*PPAR-δ*)^5^. We identified the best-hit genes that were most affected in promoter methylation levels by CRF across the four organs, namely *Acot5-ps1*, *LOC102547081*, *Stk24*, *Tuba4a*, *Sfmbt2*, and *LOC100359655*. All these best-hit genes were observed in the soleus muscle. These genes may be involved in intracellular signaling (*Stk24*), structural formation (*Tuba4a*), and gene regulation and developmental processes (*Sfmbt2*). However, some of these genes have been poorly characterized. In addition, an organ-specific comparison of promoter methylation levels across four organs showed that the number of genes with *p* < 0.05 was highest in the soleus muscle with 1427 genes. Based on our unique analysis, these results suggest that the soleus muscle methylome is more strongly and extensively affected by CRF than other organs known to be affected by exercise, such as the hippocampus, heart, and large intestine. We also identified a set of seven common genes that exhibited differences in promoter methylation levels across all four organs between the LCR and HCR groups. Some of these genes have been implicated in cell regulation, including metabolism (*Stk24*), RNA processing (*Ddx46*), and cell adhesion (*Dsc3*). Conversely, pseudogenes (*Klf5-ps2*) and several genes followed by a Gene ID numbers after “LOC” are largely undefined in their physiological roles and specific functions in cellular processes. It is well known that exercise promotes systemic health benefits through adaptations in various organs, such as the brain, heart, lungs, and liver^11,43,44^, and exercise is one of the promising “geroprotectors” in humans to date^45^. The mechanisms underlying the healthspan-promoting effects of exercise are not fully understood; however, changes in gene expression patterns due to alterations in global and gene-specific promoter methylation may partially explain this physiological phenomenon^38^. In addition to the fact that promoter methylation is most affected in the soleus muscle, the fact that the promoter methylation of seven genes is commonly affected by CRF in various organs may have important implications for understanding the molecular basis of CRF and longevity. Further studies are necessary to elucidate organ-specific and/or organ-common regulatory mechanisms of DNA methylation patterns and their interorgan interactions in exercise adaptation through exercise training interventions.

In addition, we identified organ-specific genes whose methylation levels were affected by CRF. The genes identified in the heart were *LOC100911237*, *Clec2d*, *Stk24*, *Rara*, *Mansc4*, and *Zdhhc7*, which intersect complex cellular networks and are involved in immune responses (*Clec2d*), cell signaling and stress responses (*Stk24*), gene regulation and development (*Rara*), potential protein modifications (*Mansc4* and *Zdhhc7*), and other cellular activities. The hippocampus-specific genes identified were *hist1h2ail2*, *LOC100910200*, *LOC120095374*, *Kcnt2*, *Xrcc5*, and *Pth2*. These genes contribute to basic cellular processes, such as DNA regulation (*Hist1h2ail2*), repair (*Xrcc5*), ion channel regulation (*Kcnt2*), hormone regulation (*Pth2*), and chromatin organization (*Hist1h2ail2*). The large intestine-specific genes identified were *Klk5l*, *LOC120103016*, *Nup42*, *Hibadh*, *H2aj*, and *Stk24*. These genes may be involved in protease activity (*Klk5l*), nuclear transport (*Nup42*), lipid metabolism (*Hibadh*), chromatin structure (*H2aj*), and intracellular signal transduction (*Stk24*). However, several of these genes have been poorly characterized. The soleus muscle-specific genes and their physiological roles have been described above. Organ specificity in the gene-specific promoter methylation status associated with CRF emphasizes the need to consider the unique characteristics of each organ when investigating the effects of CRF on epigenetic alterations.

Notably, exhaustive statistical analysis revealed that epigenetic age, epigenetic age acceleration, GMM, and MME exhibited different signatures among the four organs targeted in this study. More specifically, both epigenetic age and epigenetic age acceleration values were highest in the large intestine and the values of the intersection clock were highest in the large intestine, followed by the soleus muscle. GMM and MME exhibited higher values in the large intestine than in the heart, hippocampus, and soleus muscle. Moreover, GMM and MME values were second highest in the heart. Furthermore, epigenetic age acceleration was not associated across organs within the same individual. Recent results from animal studies using proteomics and transcriptomics suggest that the rate of aging varies not only between individuals but also between organs within individuals^46,47^, consistent with our results on organ distinctions in epigenetic aging. Similar studies have been reported in humans^48^, and part of the multi-organ aging network has also been elucidated, in which biological aging in specific organs influences the progression of aging in other organs^49^. On the other hand, epigenomics-based studies suggest that age-related DNA methylation changes are organ-specific in rodents^23,50^ and human^51^, but the differences in epigenetic age acceleration among organs are largely unknown. Therefore, our findings provide important insights into the biology of aging and emphasize the need to validate rejuvenation strategies in the context of the organ-specific nature of epigenetic aging. Further evidence is required to elucidate the organ-specific effects of exercise on organ-specific epigenetic aging.

In conclusion, while the applied blood-based rat epigenetic clocks do not consistently reflect the differences in CRF in any organ, higher CRF is associated with a younger state according to GMM and MME. Our results also indicate that CRF regulates promoter methylation of various genes in an organ-specific and organ-common manner. We also demonstrated that epigenetic aging exhibits different signatures in different organs and that they are not consistent across organs. These findings emphasize the potential involvement of CRF in organ-specific epigenetic aging and gene-specific regulation of promoter methylation, providing novel insights into the complicated interplay between CRF, epigenetic regulation, and aging processes.

## 4. Methods

### 4.1. Experimental animals

Sixteen female LCR and HCR rats of the 44th generation (23–24 months old) were used in this study. LCR and HCR rat models were obtained by artificially selecting low- and high-running capacities from genetically heterogeneous N:NIH stock rats^14^. Rats were transported by air from The University of Toledo (Toledo, OH, USA) to the Hungarian University of Sport Sciences (Budapest, Hungary). The rats were housed in temperature-controlled animal rooms maintained on a 12:12-h light-dark cycle. The animals were fed *ad libitum* on a standard chow diet and water. This study was conducted in accordance with the requirements of The Guiding Principles for Care and Use of Animals, EU, and was approved by The National Animal Research Ethical Committee (Hungary) (PE/EA/62-2/2021).

### 4.2. Maximal oxygen uptake test

All rats were acclimated to a motor-driven animal treadmill in a closed chamber customized for rats (Columbus Instruments, USA) prior to VO_2max_ measurements. Briefly, the animals rested on a running belt for 5 min, and the running speed was gently increased. Acclimatization runs were performed for 5–10 min at speeds of 5–25 m/min for two consecutive days. VO_2max_ was measured in the HCR/LCR rats by beginning the run at a speed of 5 m/min after 5 min of rest and then increasing the speed by 5 m/min every 2 min until exhaustion. The VO_2max_ was determined for each animal using the following criteria: (i) no change in VO_2_ with increasing speed, (ii) rats could no longer maintain their posture on the treadmill, and (iii) the respiratory quotient (RQ = VCO_2_/VO_2_) > 1^52,53^. More specifically, VO_2max_ values were recorded when the animal met one of the three criteria. We also calculated the total running time and total running distance at the time point of VO_2max_ value onset. Notably, VO_2max_ measurements were carried out in an infrared light-dark environment, at a temperature of 22 ± 2 °C and a humidity of approximately 55 ± 10 %.

### 4.3. Organ removal and DNA extractions

After completing the VO_2max_ measurements, dissection was performed after at least 48 h of rest. More specifically, rats were deeply anesthetized by intraperitoneal injection of a ketamine (Richter, concentration: 100 mg/mL) /xylazine (Produlab Pharma, concentration: 20 mg/mL) cocktail at a dose of 0.1 mL/10 g body weight and intraperitoneally perfused with heparinized ice-cold saline. The hippocampus, heart, soleus muscle, and large intestine were removed when the rats did not respond to tail compression. Organ samples were stored at −80 °C until DNA extraction. One whole side of the hippocampus, left side of the soleus muscle, and middle part of the large intestine, which corresponds to the colon, were used for further analysis. Whole hearts were pre-homogenized in 2× volume of Phosphate-buffered saline for each organ weight. DNA was extracted using the PureLink Genomic DNA Mini Kit (Thermo Fisher Scientific, USA) according to the manufacturer’s instructions. The extracted DNA was dissolved in 50 μL of PureLink Genomic Elution Buffer (10 mM Tris-HCl, pH 9.0, 0.1 mM ethylenediaminetetraacetic acid. Prior to DNA methylation measurement, DNA samples were adjusted to a concentration of ≥35 ng/μL with a purity of A260/280 >1.7.

### 4.4. Global methylation measurement by RRBS

RRBS libraries were generated using the Premium RRBS Kit V2 (Diagenode, Seraing, Belgium) as described previously^54^. Briefly, 100 ng of isolated DNA was digested with the MspI restriction enzyme, followed by end repair, adaptor ligation, and size selection using Ampure beads. Each library was quantified by quantitative PCR to determine the library concentration, and eight samples were pooled in equimolar amounts using an Excel pooling tool provided by the manufacturer. The pooled samples were subjected to bisulfite treatment, purification, and PCR amplification, according to the manufacturer’s instructions. Library concentrations were quantified using the Qubit dsDNA HS Assay (Life Technologies) and library size distribution was measured using a Bioanalyzer High-Sensitivity DNA chip. The libraries were sequenced by multiplexing eight libraries per lane on an Illumina HiSeq2500 sequencer in 1 × 50 single-end mode. Sequencing reads were trimmed using CutAdapt to remove adapter sequences. After read trimming, bisulfite alignment to the mRatBN7.2 (GCA_015227675.2) reference sequence and methylation calling were performed using Bismark v0.24.129. We filtered the aligned data to retain only cytosines with > 5× coverage in at least 90 % of samples. Methylation values for samples at CpG sites with < 5× coverage were set to Not Available. There were 1 327 821 such highly covered and mostly common CpG sites.

### 4.5. Development of RRBS-based rat epigenetic clocks

We developed three RRBS-based rat clocks using an available dataset from the GSE161141 dataset^17^. First, we downloaded the processed methylation files (CGmaps) for the 134 whole blood samples of male rats (Rattus norvegicus, F344 strain). Then we lifted the genomic positions from the rn6 to the rn7 genomes using the *liftover* Python package. For further analysis, we used only the CpG sites that were covered by at least five reads in at least 90 % of the samples (there were 1 028 452 such highly covered and mostly common CpG sites).

Rat clock 1: We filled in the missing values using the mean methylation values of the whole data, then trained glmnet (ElasticNet with alpha = 0.5) at 80 % and tested it on 20 % of the samples (**Fig.S1A**). The performance of the test set was mean absolute error (MAE) = 4.3 months and *r* = 0.88 (**Fig.S1A**). The clock used 39 CpG sites in total. We applied the clock (Rat clock 1) on our RRBS rat data after selecting the data for highly covered methylation values (≥ 5 reads) and filling missing values by the mean methylation values of the covered clock CpG sites.

Rat clock 2: First, we found common CpG sites in the two methylation tables (the Levine et al. [2020] data set^17^ and our rat data set). In total, 307 337 common CpG sites were identified. We then trained *glmnet* on 80 % of the samples from the Levine et al. (2020) dataset and tested it on 20 % using only 307 337 common CpG sites (**Fig.S1B**). The performance of the test set was: MAE = 4.4 months and *r* = 0.87 (**Fig.S1B**). The clock used 27 CpG sites in total. We applied this clock (Rat clock 2) to our rat data, which were restricted to common CpG sites. We filled missing values by the mean methylation values of the covered clock CpG sites. Given the number of samples fulfilling the number of CpG sites used for each Rat clock (39 and 27 sites, respectively), Rat clock 2 had a higher prediction reliability than Rat clock 1. Age acceleration was calculated for the two rat clocks. In the age acceleration plots, we considered only highly covered samples (when at least 75 % of the clock CpG sites were covered).

Rat intersection clock: We applied our new epigenetic clock method^18^ to 64 rat samples. We trained the intersection clock on the Levine et al. (2020) dataset to obtain a rat clock. Details of the intersection clock are provided in the Methods section of the original study. In this section, we describe it, here, briefly. The goal of this approach was to maximize the use of CpG sites in the training and test sets. For each test sample, we determined the intersection of CpG sites between the training and test datasets. Subsequently, we restricted the training set and test sample to the intersected CpG sites and used the restricted training set and test sample to train and test clocks by 5-fold cross-validation using ElasticNet (glmnet) (**Fig.S1C**). The mean prediction (MeanPred) of the five final models was used to predict the age of the restricted test sample. In this study, we considered only test samples with at least 100 000 intersecting CpG sites.

### 4.6. Global methylation analysis (GMM and MME)

We calculated the GMM for each sample as the average methylation level. Furthermore, we calculated MME for each sample as the average methylation entropy. Methylation entropy^41^ was calculated using the Shannon entropy of each individual CpG site of a sample:

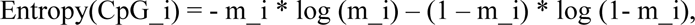

where m_i is the methylation level of the CpG site and log is the 2-based logarithm.

### 4.7. Promoter analysis of individual genes

We used the methylation table of the 64 rat samples described above (containing only 1 327 821 CpG sites covered by at least five reads in at least 90 % of the samples). Promoter region of each gene was determined as [−1500, +500] bp from transcription start site following the direction of transcription. We used the annotation file GCF_015227675.2_mRatBN7.2_genomic.gtf downloaded from the National Center for Biotechnology Information File Transfer Protocol repository (https://ftp.ncbi.nlm.nih.gov/genomes/all/GCF/015/227/675/GCF_015227675.2_mRatBN7.2/). The mean methylation levels of the CpG sites in the promoter region of each sample were calculated.

### 4.8. Statistical analysis

Correlations were evaluated by Pearson correlation coefficient (‘r’) and the corresponding two-sided *p*-values using the stats.pearsonr function of the python package scipy (stats module). We used a two-sided *p*-value in the study; statistical significance was set at *p* < 0.05. If *p*-values were indicated by an asterisk, we used the following notations: ns, *p* > 0.05; *, 0.01 < *p* ≤ 0.05; **, 0.001 < *p* ≤ 0.01; ***, *p* ≤ 0.001; and ****, *p* ≤ 1.00e-04.

## Acknowledgements

We would like to thank The University of Toledo, lab manager Samantha J. McKee, M.S. for expert phenotyping, care, and maintenance of the LCR/HCR selectively bred rat colony, and Dr. Lisa J. Root and Ashley Kurth from the Department of Laboratory Animal Resources for the attention required for the careful export of the aged LCR and HCR rats.

## Author contributions

Conceptualization, T.K. and Z.R. Methodology, T.K., C.K. and Z.R. Investigation, T.K., C.K., F.T., Z.B., L.Z., P.B., A.K., E.K. and Z.R. Statistical analyses and creating the figures, C.K. Writing – Original Draft, T.K, C.K. and Z.R., Writing – Review & Editing, T.K., C.K., L.B., L.G.K., S.L.B. and Z.R., Supervision, Z.R.

## Non-standard Abbreviations and Acronyms

CRF: cardiorespiratory fitness
GMM: global mean DNA methylation
HCR: high-capacity runner
LCR: low-capacity runner
MAE: mean absolute error
MME: mean methylation entropy
RRBS: reduced representation of bisulfite sequencing
VO_2max_: maximal oxygen uptake.

---

**Figure S1.**
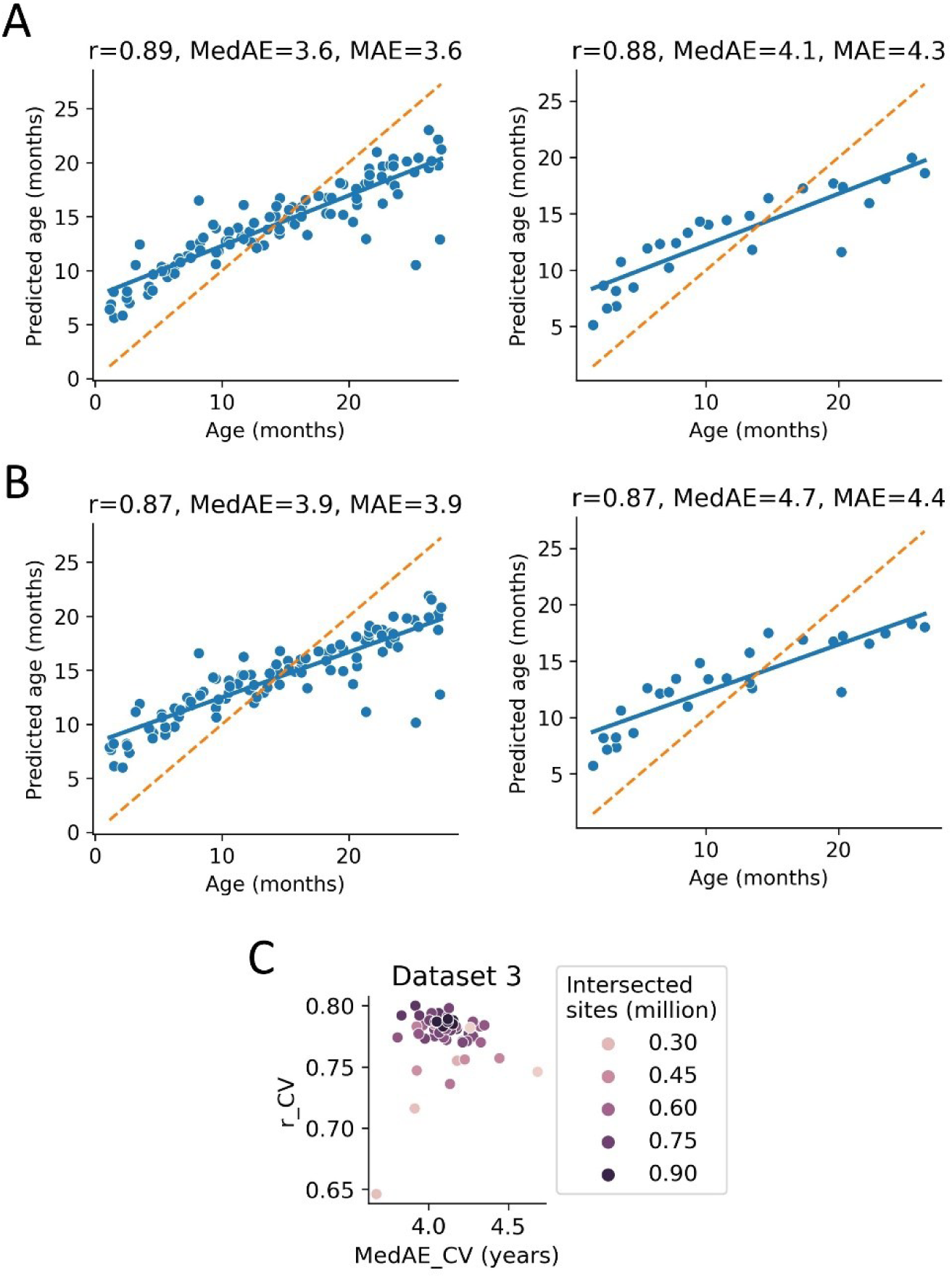
Training and testing of the Levine rat blood dataset. **A.** Training and testing of the Rat clock 1. **B.** Training and testing of the Rat clock 2. **C.** Number of intersected CpG sites and cross-validation (CV) model performance, assessed by Pearson correlation coefficient (*r*) and median absolute error (MedAE), of the intersection clock.

**Figure S2.**
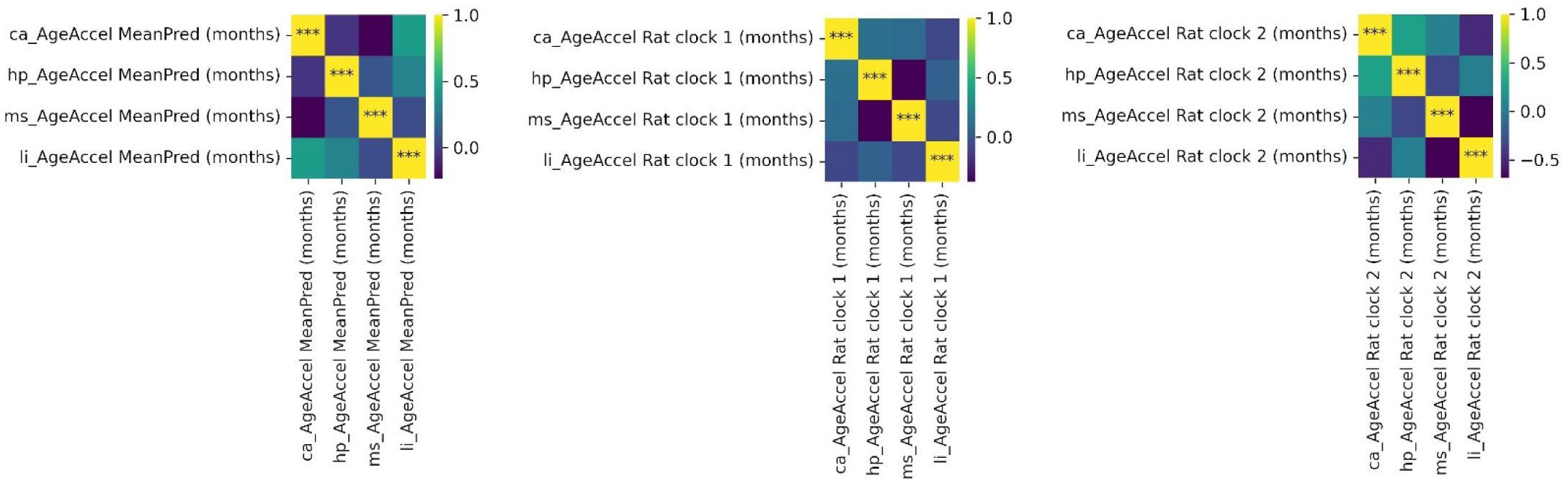
Inconsistency of epigenetic age acceleration across organs in the same individual. We calculated the Pearson correlation coefficient between age accelerations of organ-pairs. The three plots are linked to the epigenetic age acceleration based on the three rat clocks (intersection clock, Rat clock 1, and Rat clock 2, respectively). ca: heart, hp: hippocampus, ms: soleus muscle, li: large intestine. *: 0.01 < *p* ≤ 0.05, **: 0.001 < *p* ≤ 0.01, ***: *p* ≤ 0.001, ****: *p* ≤ 0.0001, ns: not significant.

